# Avian influenza virus circulation and immunity in a wild urban duck population prior to and during a highly pathogenic H5N1 outbreak

**DOI:** 10.1101/2024.02.22.581693

**Authors:** Jordan Wight, Ishraq Rahman, Hannah L. Wallace, Joshua T. Cunningham, Sheena Roul, Gregory J. Robertson, Rodney S. Russell, Wanhong Xu, Dmytro Zhmendak, Tamiru N. Alkie, Yohannes Berhane, Kathryn E. Hargan, Andrew S. Lang

## Abstract

Highly pathogenic avian influenza (HPAI) H5N1 clade 2.3.4.4b viruses were first detected in St. John’s, Newfoundland, Canada in late 2021, with the virus rapidly spreading across the western hemisphere over the next year. To investigate the patterns of avian influenza virus (AIV) infection and immune responses subsequent to the arrival of H5N1, we sampled the wild urban duck population in St. John’s for a period of 16 months after the start of the outbreak and compared these findings to archived samples. Antibody seroprevalence was relatively stable before the outbreak (2011-2014) at 27.6% and 3.9% for anti-AIV (i.e., NP) and H5-specific antibodies, respectively. During the winter of 2022, AIV-NP and H5-specific antibody seroprevalence both reached 100%, signifying a population-wide infection event. As expected, population-level immunity waned over time, and we found that ducks were seropositive for anti-AIV-NP antibodies for around twice as long as for H5-specific antibodies. The population was H5 seronegative to the latter approximately six months after the initial H5N1 incursion. In late February 2023, H5N1 clade 2.3.4.4b viruses were again detected in the duck population as a result of a second incursion into Newfoundland from Eurasia, which resulted in a second population-wide infection event. We observed a clear relationship of increasing antibody levels with decreasing viral RNA loads that allowed for interpretation of the course of infection and immune response in infected individuals and applied these findings to two cases of resampled ducks to infer infection history. Our study highlights the significance of applying both AIV surveillance and seroprevalence monitoring to provide a better understanding of AIV dynamics in wild populations, which may be crucial following the arrival of 2.3.4.4b H5Nx subtypes to assess the threats they pose to both wild and domestic animals, and to humans.

## Introduction

Wild birds are the reservoir hosts of avian influenza viruses (AIVs), with waterfowl being one of the main reservoir groups and vectors by which AIVs are spread, along with gulls, shorebirds, and seabirds [1–4]. AIVs are classified as low pathogenic (LPAIV) and highly pathogenic (HPAIV) based on their virulence and the patterns of mortality they cause in chickens [5]. LPAIV infection of waterfowl rarely results in overt disease symptoms, with birds usually recovering within a matter of days. Dabbling ducks (Anatinae) infected with HPAIV H5Nx subtypes, similar to LPAIV infection, can be minimally affected while shedding large quantities of virus, with mild disease symptoms and delayed local movements in some cases [6–9]. While many species of diving ducks (Aythyinae) also appear to be minimally affected, some such as tufted ducks (*Aythya fuligula*) have been shown to be particularly prone to experience symptomatic HPAIV H5Nx infections and can exhibit severe infection outcomes and high rates of mortality [7,10,11]. Recently, mortality in dabbling ducks due to HPAIV infections has been observed, representing a new pattern for HPAIV dynamics in one of the main reservoir hosts [11–13].

HPAIV clade 2.3.4.4 H5Nx viruses have been circulating with increasing frequency in wild birds in Eurasia and Africa since 2005 [10,14–16], with the first incursion of an A/goose/Guangdong/1/1996 (Gs/GD) lineage H5N8 clade 2.3.4.4 viruses into North America taking place in 2014 [17]. This virus and a reassortant H5N2 virus did not persist and become established in North American wild bird populations. However, new incursions of clade 2.3.4.4b viruses starting in late 2021 have resulted in the extensive reassortment with North American lineage LPAIVs, widespread circulation of H5Nx viruses throughout North and South America within a wide array of avian hosts, and multiple spillover events into mammals [18–21]. These HPAIVs now seem to be part of the endemic viral population in wild birds globally.

AIV surveillance in wild birds has been a global focus for decades, with the aim of understanding viral dynamics and identifying circulating strains in different regions and species. However, this has not been without challenges. Non-gallinaceous birds infected with LPAIVs are usually asymptomatic and test positive for viral RNA for only a very short period, generally 5-11 days with considerable variation by species, body condition, exposure dose, viral strain(s), and infection history [22–26]. This provides a very narrow sampling window for detection of active infections, meaning there are certainly infections and outbreaks that go undetected. An increasing number of serological studies have helped address this shortcoming, by which past AIV infection can be documented via detection of anti-AIV antibodies in the peripheral circulation for a period of months [25,27–29]. A combined approach of AIV infection surveillance and serology can therefore help capture AIV dynamics over a longer time frame, allowing interpretation of both active and past infections in populations [9,30,31].

Several groups have shown that antibody levels decrease over time following infection of AIV-naïve captive ducks with a variety of different AIVs, as expected. However, after homologous and heterologous challenge, antibody levels rebound in a matter of days and the ducks are often protected from clinical disease [24,25,32]. Unfortunately, the majority of these studies, including all those on HPAIVs [23,29,33–35], have been performed at timescales of weeks, and therefore the duration of protection from subsequent re-infection by HPAIVs is currently unknown.

A GsGd lineage clade 2.3.4.4b HPAI H5N1 virus was identified in a great black-backed gull (GBBG, *Larus marinus*) that died in November 2021 in St. John’s, Newfoundland and Labrador, Canada, and was found to be closely related to viruses circulating in northwestern Europe in the spring of 2021 [36]. Shortly after this first detection, the virus was identified in an exhibition farm in the area that housed primarily domestic fowl, which resulted in mass mortality [36]. Sampling of wild urban ducks within the area began about a week later and active H5N1 infection was detected in the duck population in late December 2021. There was no observed or reported morbidity or mortality of waterfowl in the St. John’s area between November 2021 and January 2023 [37].

The island of Newfoundland lies off the eastern coast of mainland North America and is within the extreme eastern edge of the Atlantic Americas Flyway [3]. AIV dynamics and subtype diversity in ducks, gulls, and seabirds have been studied in the region since 2007 to understand possible linkages with European strains, and this work included some focus on the wild urban duck population in the St. John’s area [38,39]. A broad diversity of AIV subtypes with high rates of strain turnover have been detected in this population, including an H5 virus (A/American black duck/Newfoundland/1181/2009(H5N4)) [38], but no HPAIVs had been detected in the province prior to late 2021. The dynamics of duck movement in this area are well understood through banding programs and associated recaptures and resightings, which have shown this population of ducks is largely composed of non-migratory individuals [38]. Their use of public urban waterbodies means that many are accustomed to humans, allowing capture and recapture or resighting of the same individuals multiple times throughout the year.

The aim of this study was to thoroughly investigate patterns of AIV infection and immunity in this duck population over a period of approximately 16 months after the first arrival of GsGd lineage H5N1 to North America in November 2021. We accomplished this goal by employing a combination of AIV surveillance to understand when infection was occurring and serology, specifically of general anti-AIV-NP as well as H5-specific antibodies, to understand immune responses. This work focused on how immunity changed over time while epidemiological information provided context and timing of infection(s) and bird movements.

## Materials and Methods

### Ethics statement

This work was carried out under the guidelines specified by the Canadian Council on Animal Care with approved protocols 11-01-AL, 12-01-AL, 13-01-AL, 14-01-AL, 17-05-AL, and 20-05-AL from Memorial University’s Institutional Animal Care Committee, biosafety permit S-103 from Memorial University’s Institutional Biosafety Committee, and banding and sampling operations under Federal Bird Banding and Scientific Research Permit 10559.

### Bird capture and sampling

Wild ducks were caught either by hand or bait trapping at several locations in or near the city of St. John’s, Newfoundland, including Bowring Park (47.528862°, −52.745943°), Commonwealth Pond (47.500765°, −52.789646°), Kenny’s Pond (47.591366°, −52.715759°), Kent’s Pond (47.589212°, −52.722767°), Mundy Pond (47.551419°, −52.741791°), Quidi Vidi Lake (47.579076°, −52.699627°), and Topsail Pond (47.524388°, −52.903371°). Sampling occurred in the fall and early winter months from 2011 to 2014, and at 11 timepoints through 2022 and 2023 during the ongoing HPAI outbreaks (see Supplementary Data). Bird age was determined using plumage aspect and cloacal characteristics [40,41]. Age categories included hatch year (HY), after hatch year (AHY), second year (SY), and after second year (ASY). Hatch year birds that have not yet fledged are denoted as local (L). All birds were banded with a metal leg band issued by the Canadian Wildlife Service Bird Banding Office.

Capture efforts targeted primarily mallards (MALL, *Anas platyrhynchos*), American black ducks (ABDU, *A. rubripes*), northern pintails (NOPI, *A. acuta*), and occasionally hybrid ducks that were a combination of ABDU/MALL/feral domesticated ducks (*Anas* spp.). Additional species were sampled opportunistically in 2022 and 2023, specifically American wigeon (AMWI, *Mareca americana*), Eurasian wigeon (EUWI, *M. penelope*), and lesser scaup (LESC, *Aythya affinis*). As there were a limited number of ducks that were AIV RNA-positive at the time of capture, we included AIV surveillance and serology data from several seabird species originating from other work to explore a larger dataset for an analysis on the relationship between RNA load and antibody levels. These species were also impacted by outbreaks of HPAI H5N1 that occurred during the summer of 2022. Data for 100 seabirds were included, with Atlantic puffins (ATPU, *Fratercula arctica*), black-legged kittiwakes (BLKI, *Rissa tridactyla*), and common murres (COMU, *Uria aalge*) from Gull Island (47.262509°, −52.773526°) sampled between June and August 2022 and in June 2023, and northern gannets (NOGA, *Morus bassanus*) from Cape St. Mary’s (46.818668°, −54.182652°) sampled in July 2022.

### Observations of wild bird movements

Observations and remarks regarding the patterns of arrival of migratory individuals and timeline and movements of non-resident species were primarily made directly while working in the field, with additional support provided by local experts, other birders in the region, sightings posted to birding social media pages, and submissions to ebird.org [42].

### Bird banding and encounter data

To provide further support of this wild urban duck population being comprised of primarily resident individuals, we obtained banding and encounter data (reporting of a bird band) for all dabbling and diving ducks banded within a 20 km radius of St. John’s, Newfoundland from 1 January 2010 to 17 May 2023 from the Canadian Wildlife Service Bird Banding Office.

### Sampling periods

For analysis purposes, samples collected between 2011 and 2014 were grouped into sampling seasons. Sampling occurred across multiple months between September and March of each period, and are referred to as the 2011-2012, 2012-2013, 2013-2014, and 2014-2015 seasons. Samples collected after H5N1 was first detected in 2021 were grouped as follows: winter-spring 2022 (samples collected between January and May 2022), summer-fall 2022 (samples collected between July and September 2022), and then separately for samples collected in the months of February, March, and April 2023. Specific details about when each sample was collected can be found in the Supplementary Data.

### Serology

Two to three millilitres of blood were drawn from the brachial wing vein of each captured individual. Serum was separated from clotted blood by centrifugation at 3,000 𝑥 g for ten minutes and subsequently stored at −20°C for future analysis. For the 19 samples collected in January and February 2022 and for 28 archived samples, AIV competitive enzyme-linked immunosorbent assays (cELISAs) were performed at the National Centre for Foreign Animal Disease (NCFAD) laboratory as previously described [43]. Serum from 12 of these 19 samples were later re-tested using the IDEXX AI MultiS Screen Ab test (IDEXX Canada, Product # 99-12119) as per the manufacturer’s instructions, which detects antibodies against influenza A nucleoprotein (NP) [44], and this assay was used for all samples collected from March 2022 onwards and all other archived samples. A sample to negative control ratio (S/N) of < 0.5 was considered positive for influenza antibodies. As some studies have employed a S/N ratio of < 0.7 for positivity [28,44,45], this value is shown on relevant figures for comparison purposes. All archived samples collected between 2012 and 2014 were re-tested for the present study to confirm the original results. Samples from the 2011-2012 period were no longer available, therefore the previously published data were used [38]. Due to a lack of serum for 7 individuals sampled in January and February 2022, re-testing using the IDEXX assay could not be performed, therefore analyses using S/N ratios were performed for 76 individuals, instead of all 83 ducks sampled between 2022 and 2023. All sera positive for anti-NP antibodies were subsequently tested at the NCFAD for antibodies specifically against subtype H5 [46]. The two seronegative individuals sampled in February 2022 were removed from the week-by-week analysis as they were believed to have not been present at the time of the population-wide infection event (see Discussion). Additionally, the six individuals sampled on 23 August 2022 were also removed from this analysis as they were presumed to be primarily migratory individuals, coinciding with the large influx of individuals during this post-breeding migration period.

### Swab samples and RNA isolation

Oropharyngeal and cloacal swabs were collected from all individuals from 2022 to 2023 and the paired swabs were pooled into a single tube of Multitrans viral transport medium (Starplex Scientific, Product # S160-100) and represent a single sample per individual. Samples were stored in a cooler on ice and an aliquot was removed for RNA isolation within six hours, and samples were subsequently stored at −80°C. RNA was isolated from 140 μL of each sample using the Qiagen Viral RNA Mini Kit (Qiagen, Product # 52906) as per the manufacturer’s instructions and stored at −80°C until further analysis.

### Screening for influenza A viruses

Real-time RT-PCR was performed using AgPath-ID™ One-Step RT-PCR reagents (Applied Biosystems, Product # 4387424) on a StepOnePlus Real-Time PCR System (Applied Biosystems). All samples were screened for the presence of the influenza A virus (IAV) matrix gene and subsequent positives were screened for the H5 subtype of the haemagglutinin gene. RT-qPCR primers and probes, and cycling conditions were adapted from Spackman (2020) with some modifications. For the initial RT-qPCR targeting the influenza matrix gene, 25 μL reactions were prepared using 12.5 μL of 2X RT-PCR buffer, 1 μL of 25X RT-PCR enzyme mix, 0.25 μL of 20 μM F25 (5’ - AGATGAGTCTTCTAACCGAGGTCG – 3’), 0.25 μL of 20 μM R124 (5’ - TGCAAAAACATCTTCAAGTCTCTG – 3’), 0.25 μL of 20 μM R124M (5’ - TGCAAAGACACTTTCCAGTCTCTG – 3’), 0.25 μL of 6 μM double quenched probe F64P (5’ – [FAM]-TCAGGCCCC[ZEN]CTCAAAGCCGA-[IB] – 3’) (IDT Inc., Canada), 1.67 μL of AgPath Detection Enhancer (Applied Biosystems, Product # A44941), 0.83 μL of nuclease-free water, and 8 μL of RNA. Cycling was performed in standard mode, with parameters as follows: 45°C for 20 minutes, 95°C for 10 minutes, followed by 45 cycles of 95°C for 5 seconds, and 60°C for 1 minute at which time fluorescent signal was detected. A standard curve of IAV RNA as well as no-template controls were included during each run. Thresholds were determined automatically by the StepOnePlus software based on the standard curve, and this threshold was applied after manual confirmation to determine the cycle threshold (Ct) values for each sample. Samples that yielded the characteristic amplification curve and had a Ct ≤ 45 were interpreted as positive [48–50], while those that yielded the characteristic amplification curve but did not surpass the threshold were interpreted as inconclusive and denoted as having a Ct > 45.

All samples that yielded an amplification curve for IAV matrix RT-qPCR were subsequently screened for the H5 subtype using primers, probes, and cycling conditions adapted from Spackman et al., (2002) with some modifications. The 25 μL reactions were prepared using 12.5 μL of 2X RT-PCR buffer, 1 μL of 25X RT-PCR enzyme mix, 0.25 μL of 20 μM H5_1456-NA_F (5’ – ACGTATGACTATCCACAATACTCA – 3’), 0.25 μL of 20 μM H5_1456-EA_F (5’ – ACGTATGACTACCCGCAGTATTCA – 3’), 0.125 μL of 20 μM H5_1685_R (5’ – AGACCAGCTACCATGATTGC – 3’), 0.125 μL of 20 μM H5_1685M_R (5’ – AGACCAGCTATCATGATTGC – 3’), 0.25 μL of 6 μM double quenched probe H5_1637P (5’ – [FAM]-TCAACAGTG[ZEN]GCGAGTTCCCTAGCA-[IB] – 3’) (IDT Inc., Canada), 2.5 μL of nuclease-free water, and 8 μL of RNA. Cycling was performed in standard mode, with parameters as follows: 45°C for 20 minutes, 95°C for 10 minutes, followed by 45 cycles of 94°C for 10 seconds, 57°C for 40 seconds at which time fluorescent signal was detected, and 72°C for 5 seconds. Any samples that yielded the characteristic amplification curve were interpreted as positive for H5.

Samples that tested negative for H5 were assumed to represent infection with an LPAIV. For these samples, the NEB OneTaq® One-Step RT-PCR Kit (New England Biolabs, Product # E5315S) was used to target the haemagglutinin gene. 25 μL reactions were prepared using 12.5 μL of 2X OneTaq One-Step Reaction Mix, 1 μL of 2X OneTaq One-Step Enzyme Mix, 1 μL of 10 μM HA-1134F (5’ - GGRATGRTHGAYGGNTGGTAYGG – 3’), 1 μL of 10 μM Bm-NS-890R (5’ - ATATCGTCTCGTATTAGTAGAAACAAGGGTGTTTT – 3’), 1.5 μL of nuclease-free water, and 8 μL of RNA. Cycling parameters were as follows: 48°C for 60 minutes, 95°C for 5 minutes, 7 cycles of 94°C for 15 seconds, 42°C for 30 seconds, and 68°C for 3 minutes, then 35 cycles of 94°C for 15 seconds, 58°C for 30 seconds, and 68°C for 3 minutes, followed by a final extension at 68°C for 7 minutes. PCR products were subjected to electrophoresis for visualization, and amplicons were purified using AMPure XP beads (Beckman Coulter) and subjected to Sanger sequencing at The Hospital for Sick Children (Toronto, Canada).

### Classification of individual infection status

We used antibody levels (S/N ratios), AIV RNA load, and the epidemiological data, specifically the known dates of population-wide infection, to classify each individual based on their infection status. Currently infected individuals were those with detectable viral RNA. Recently infected individuals represented those that were negative for AIV RNA but were sampled within six months of the population-wide infection events or had elevated antibody levels (S/N < 0.7). They were subdivided into categories of being infected one, three, or six months previously based on the epidemiological patterns, or recently infected if the time between infection and sampling was unknown. Individuals having low antibody levels (S/N > 0.7) were classified as being either naïve or having antibody levels that had waned over time.

### Statistical analysis and data visualization

RStudio v4.1.0 [52] was used to perform data manipulation, statistical analysis, and data visualization using the packages cowplot v1.1.1 [53], data.table v1.14.2 [54], ggplot2 v3.4.0 [55], and readxl v1.4.2 [56]. In all tests where a *p*-value was generated, *p* < 0.05 was considered as significant.

## Results

### Samples collected and used in the study

A total of 217 serum samples were collected from ducks between 2011 and 2014, with the data from 38 samples from the 2011-2012 season being previously published [38]. After the first cases of HPAIV H5N1 in the province in late 2021, 83 paired swab and serum samples were collected from ducks between January 2022 and April 2023. In total, 300 duck sera are included in this study from 298 individuals, with two ducks recaptured and resampled in 2023. Paired swab and serum samples from 100 seabirds that were sampled during the summers of 2022 and 2023 were also included for an analysis on the relationship between viral RNA load and antibody levels, providing a larger dataset than available solely from the ducks.

### Movement patterns of banded ducks

In total, 1,045 ducks were banded within the St. John’s area (20 km radius) between 1 January 2010 to 17 May 2023. Of these banded birds, 176 were reported as being encountered at least once, with 172 (97.7%) being reported in St. John’s, two reported elsewhere in Newfoundland, one reported in Labrador, and one reported in Nova Scotia. Of all 176 encounters, 104 were reported dead/by hunters, 70 by recapture or resightings, and two were unspecified. This provides additional support that the wild urban duck population comprises primarily resident individuals that spend their entire lives in the same local region.

### Changes in population seroprevalence over time

Before the incursion of HPAIV H5N1 into the region in November 2021, the overall mean AIV-NP seroprevalence was 27.6% (range 17.6-52.6%) for the sampling seasons of 2011-2015. Antibodies specifically against H5 were markedly lower at a mean seroprevalence of 3.9% (range 2.2-5.6%) between 2012 and 2014 (**Fig 1**), which reflects the fact that there were no HPAI H5Nx viruses circulating in this region during this time period and that LPAIV H5 strains circulate in this population at low prevalence [38]. In the winter-spring period of 2022, one to four months after the arrival of HPAI H5N1, AIV-NP and H5-specific seroprevalence reached 90.9% (20/22) and 81.8% (18/22), respectively. This indicates that after the introduction of H5N1 to the region, most of the population was infected with this virus. During the summer-fall period of 2022, seroprevalence decreased to 45.8% (11/24) and 8.3% (2/24) for AIV-NP and H5-specific antibodies, respectively. Therefore, H5 seropositivity essentially returned to the baseline levels observed before the arrival of H5N1. In February 2023, just over one year after the original incursion, AIV-NP seroprevalence was approaching similar levels as observed over 2011-2014 at 42.9% (9/21), while H5-specific seroprevalence had increased to 19% (4/21), likely due to the beginning of circulation of an H5 virus again in the population. Approximately three weeks later in March 2023, AIV-NP seroprevalence rose to 100% (10/10) and H5 RNA was detected in four (40%) of these ducks. Only one of the ducks (10%) was seropositive for H5-specific antibodies at this time. Seven weeks later, at the end of April 2023, all six individuals sampled were seropositive for both AIV-NP and H5-specific antibodies (**Fig 1**).

**Fig 1.**
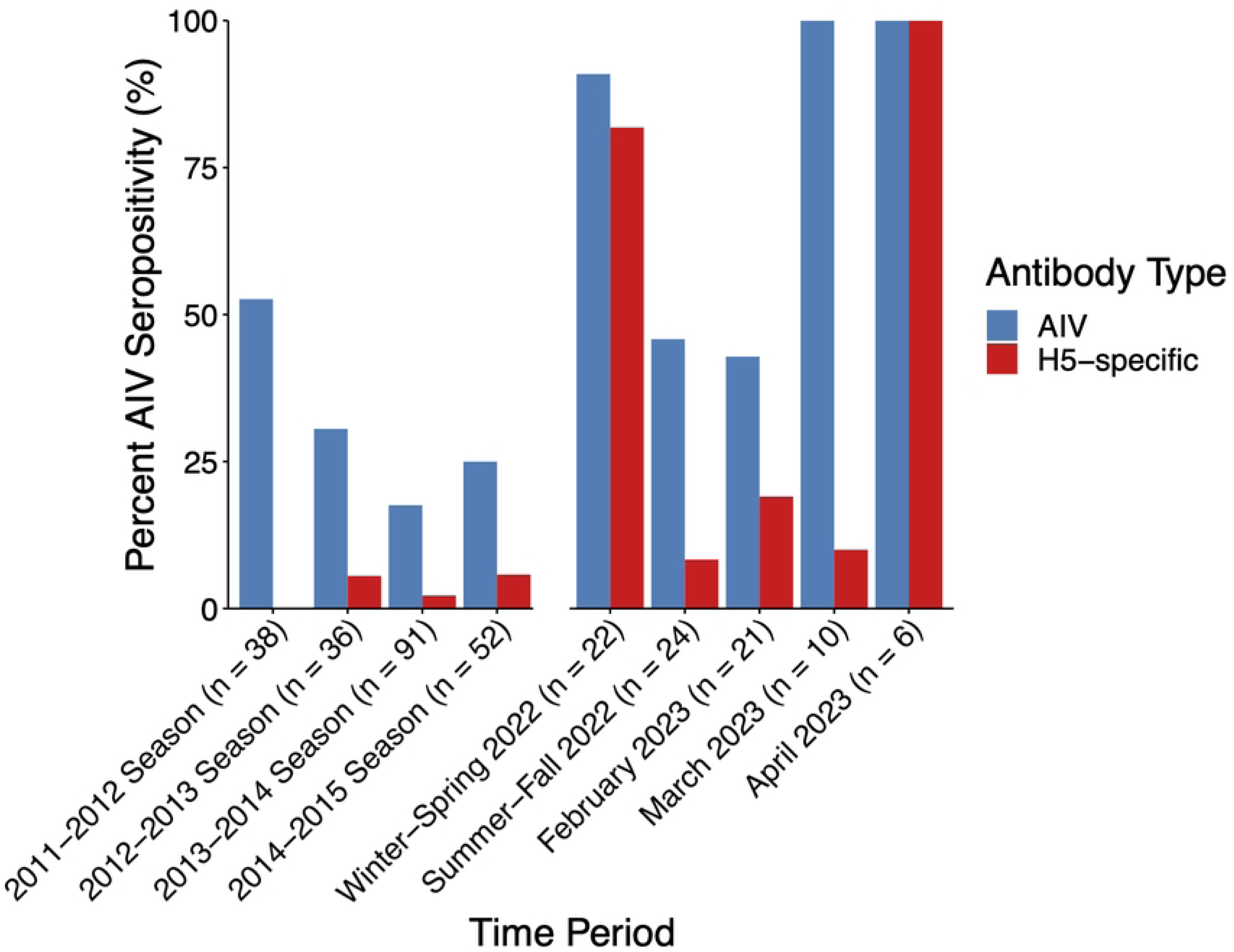
AIV and H5-specific seropositivity in the urban duck population over time. Sampling occurred in the fall and early winter months between 2011 to 2014, and then at 11 timepoints through 2022-2023 during the ongoing HPAIV H5N1 outbreak. AIV seropositivity data for samples from 2011-2012 were previously published [38] and the H5-specific ELISA was not performed with these since they were no longer available. Only samples that were positive for AIV antibodies were tested for H5-specific antibodies.

To further understand how immunity in the population changed over time, we investigated the seroprevalence (*n* = 75) over 64 weeks, specifically from 28 January 2022 through 25 April 2023, and also tested for AIV infection over this period (**Fig 2**). After the initial incursion of the GsGd lineage H5N1 virus and the population-wide infection event, seroprevalence decreased substantially over time. By approximately six months later, all ducks were seronegative for H5-specific antibodies, while half were still anti-AIV-NP antibody seropositive. AIV-NP antibodies were elevated for roughly twice as long as H5-specific antibodies (𝜒^2^= 4.97, df = 9, *p* = 0.0005) over this period. A change in AIV-NP seropositivity occurred between weeks 32 and 37, corresponding to July and August 2022, when several individuals tested positive for non-H5 AIVs (H9Nx, H11Nx, and two additional strains of unknown HA subtype). This resulted in a slight increase in seropositivity that aligned with detection of LPAIVs through the summer of 2022 to February 2023 (**Fig 2**). In March 2023, H5-subtype viral RNA was again detected, and all birds sampled were AIV-NP seropositive, with only one of these individuals seropositive for H5-specific antibodies at this time. By the end of April 2023, all individuals sampled were seropositive for both AIV-NP and H5-specific antibodies, indicating that an H5 subtype virus had again spread through the population, despite the fact that this population was infected just over a year prior. Overall, using a combination of AIV surveillance and strain subtyping, serology, and epidemiology we were able to construct a robust timeline of AIV infection and immune response in this population for the 16-month period (**Fig 3**).

**Fig 2.**
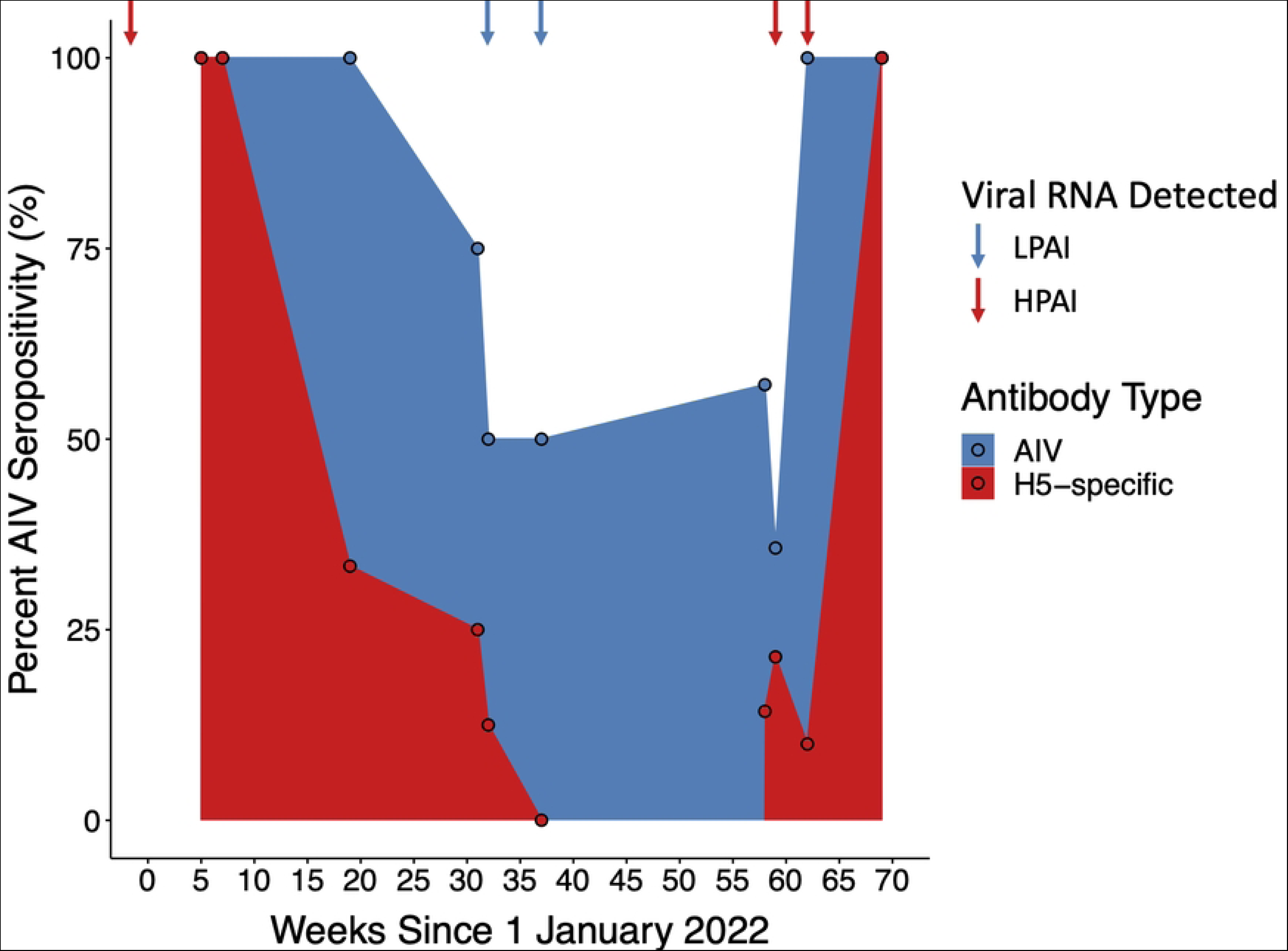
Changes in AIV and H5-specific seropositivity over the course of 16 months after the arrival of H5N1. Sampling began on 28 January 2022 and continued until 25 April 2023. Arrows above the plot correspond to AIV detections in individuals sampled at that time point, with red arrows denoting when HPAIV (H5N1) was detected and blue arrows denoting LPAIV(s) was detected.

**Fig 3.**
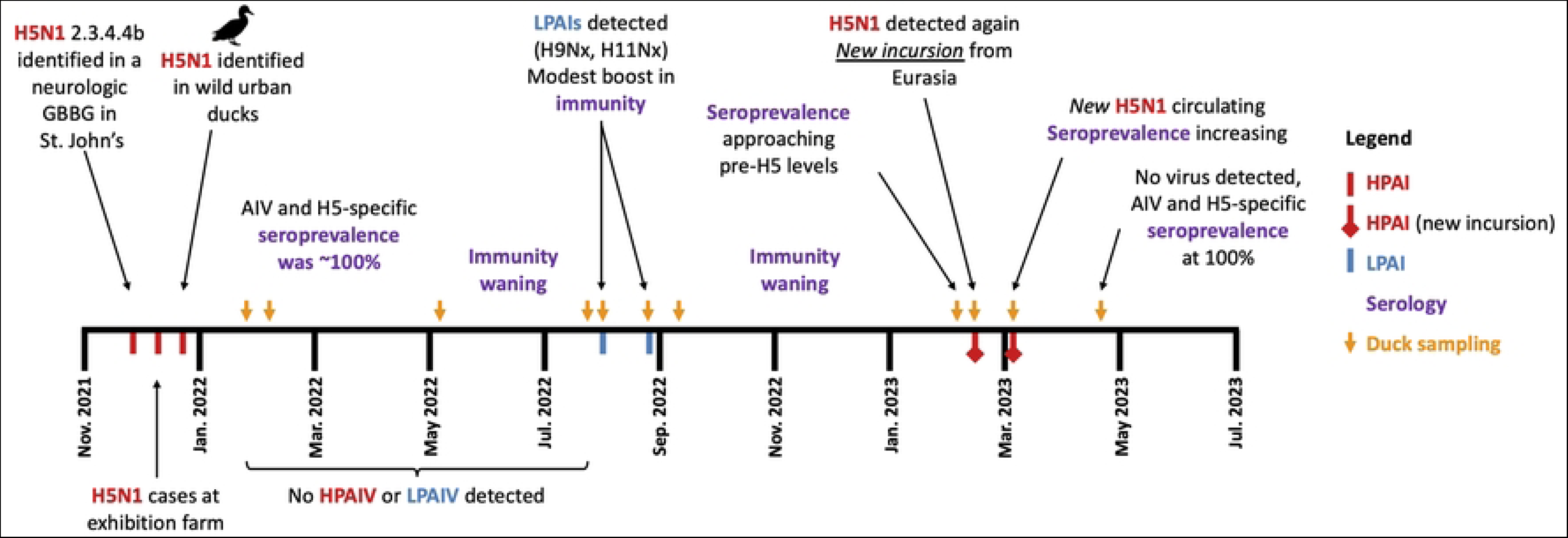
Summary of the AIV infection and immunity timeline since arrival of HPAIV H5N1 in the region.

### Immune responses in currently infected individuals

There were five individuals that were actively infected that had Ct values < 40, and these showed a negative relationship between antibody levels and viral RNA load (**Fig 4**). For ducks that were AIV RNA-negative at the time of sampling and where the time since infection was known, specifically those infected one, three, or six months prior to sampling, antibody levels decreased significantly over this time period (**Fig 4**; *F* = 5.71, df = 1,15, *p* = 0.03).

**Fig 4.**
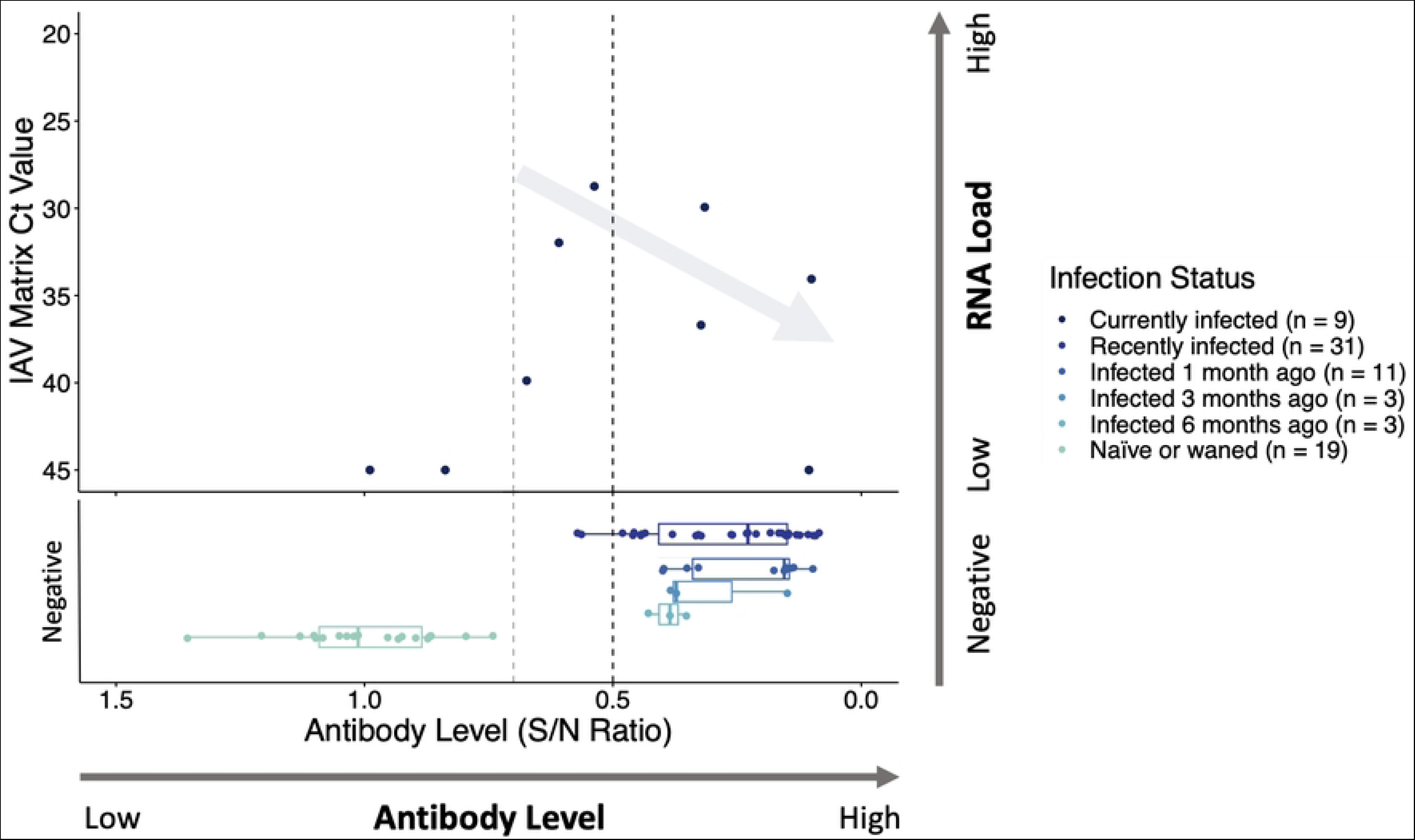
Relationship between AIV RNA load and AIV antibody levels for ducks. Each point represents an individual duck (*n* = 76), and colours indicate infection status. Individuals currently infected were those that had detectable AIV RNA loads by RT-qPCR. Samples with low RNA loads that provided expected amplification curves but that did not surpass the cycle threshold value are represented as having a Ct value of 45. Recently infected individuals were classified as such if they were AIV RNA-negative but had elevated antibody levels. For individuals sampled soon after the population-wide infection event at the start of the outbreak, these recently infected ducks are further separated as being infected one, three, or six months prior. Individuals that were seronegative and had low antibody levels (high S/N ratios) were classified as being naïve or that their antibodies had waned. An arrow denotes the expected immunological response shown by the five currently infected individuals with Ct values < 40, illustrating the observed trend of increasing antibody levels with decreasing viral RNA load. The vertical line at an S/N ratio of 0.5 represents the threshold value used to classify seropositivity, with a line at 0.7 also shown as the threshold occasionally used for this assay in other studies.

Including additional AIV prevalence and seroprevalence data obtained from various seabird species further supported the patterns observed in the ducks. Of the combined duck and seabird samples (*n* = 176), 17 individuals (9.7%) were currently infected and showed a clear relationship of increasing antibody levels with decreasing AIV viral RNA load (**Fig 5A**), an expected immunological response. A generalized additive model was used to highlight this relationship, showing the immune response in infected individuals at a population level (**Fig 5B**).

**Fig 5.**
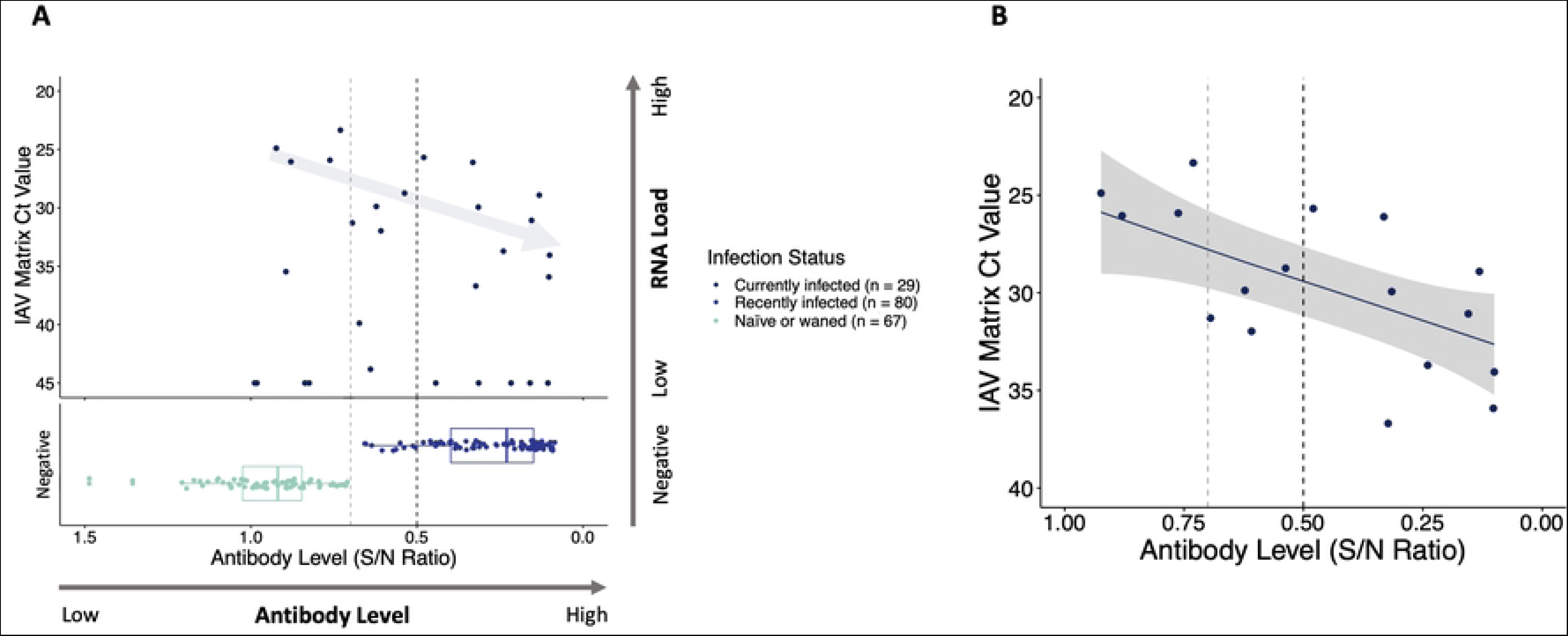
Relationship between AIV RNA load and AIV antibody levels for ducks and seabirds. Each point represents an individual and colours indicate infection status. (**A**) Data from all ducks (*n* = 76) from **Fig 4** are included in this plot, along with data for 100 seabirds (42 Atlantic puffins, 16 black-legged kittiwakes, 28 common murres, and 14 northern gannets). Individuals were classified as being currently infected, recently infected, or naïve/waned, using the same classification as in **Fig 4**. An arrow denotes the expected immunological response shown by the seventeen currently infected individuals, illustrating the relationship between increasing antibody levels with decreasing viral RNA load. These same individuals are shown in (**B**), where a generalized additive model (GAM) was fit showing the relationship between antibody level and viral RNA load, with the gray area indicating standard error of the model. The vertical lines at an S/N ratio of 0.5 represent the threshold value used to classify seropositivity, with lines at 0.7 also shown as the threshold occasionally used for this assay in other studies.

### Changes in serology of two recaptured ducks

Over the course of sampling between January 2022 and April 2023, two northern pintails were recaptured and resampled, allowing for comparison of antibody levels between the two timepoints. Both individuals were captured and recaptured at the same location, Bowring Park, and were viral RNA-negative at both sampling time points. The first, a male ASY (band # 1196-13442) was first captured on 28 January 2022 and subsequently recaptured on 7 February 2023, totalling 375 days between samples. The second, a female AHY (band # 1196-13448) was first captured on 31 July 2022 and subsequently recaptured on 7 February 2023, totalling 191 days between samples. We do not know if they were infected with AIV(s) between the sampling events, but antibody levels were lower in both individuals at the time of recapture, although to different degrees (**Fig 6**).

**Fig 6.**
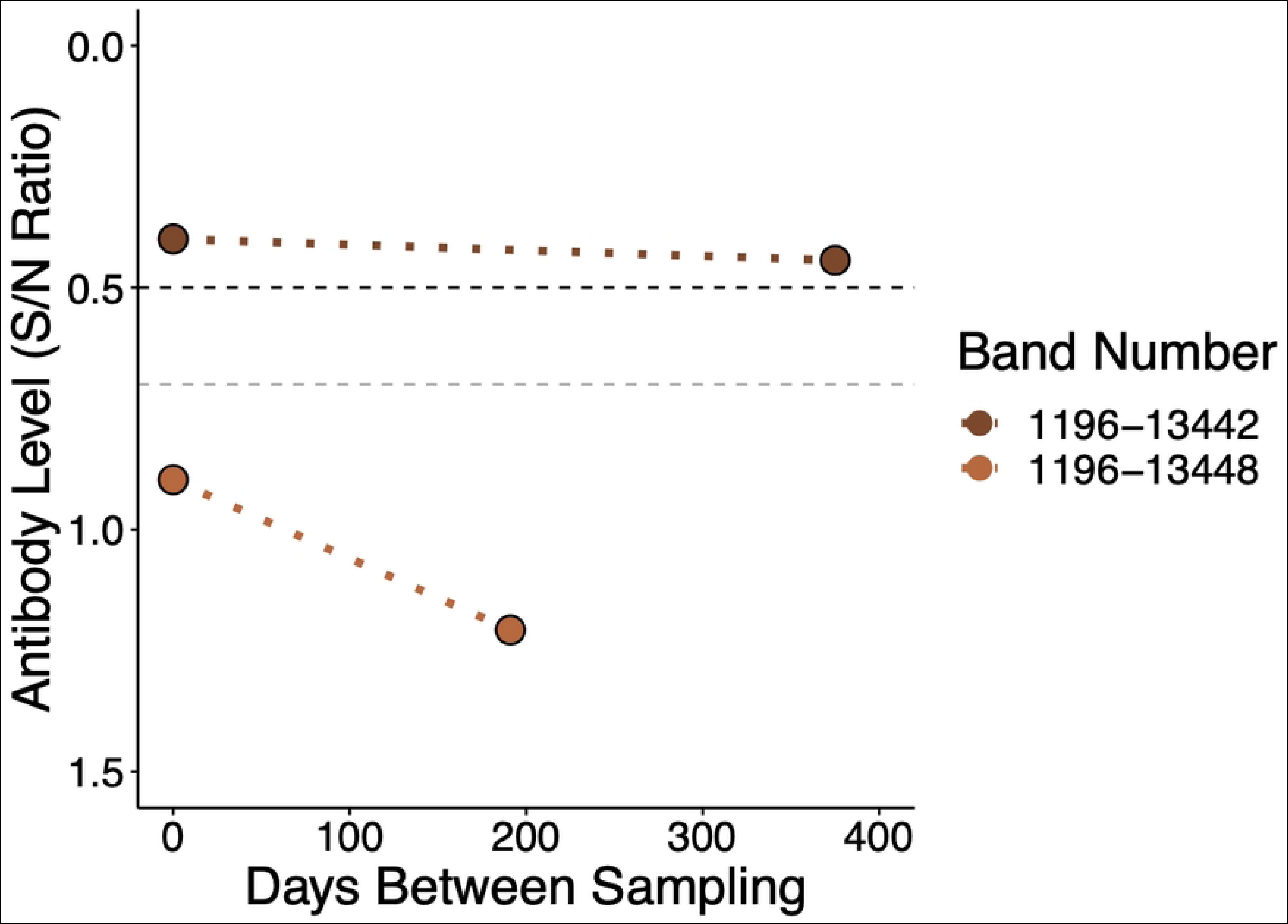
Antibody levels of two northern pintails at two timepoints. The bird with band # 1196-13442 was recaptured after 375 days and the bird with band # 1196-13448 was recaptured after 191 days. Dotted lines connect original capture and recapture values to show the change in antibody levels; these do not represent linear regressions as we do not know if or when they were reinfected with AIV between the sampling events. The vertical line at an S/N ratio of 0.5 represents the threshold value used to classify seropositivity, with a line at 0.7 also shown as the threshold occasionally used for this assay in other studies.

## Discussion

In this investigation we used a combination of AIV surveillance, serology, and epidemiology to document AIV infection and immune responses in an urban duck population before the incursion of the GsGd lineage H5N1 virus in late 2021, and over a period of 16 months through the ongoing outbreak. Through repeat sampling of the same population, we investigated changes in antibody levels through two population-wide HPAIV infection events and examined patterns of immune response over the course of AIV infection on individual scales. Using this combined information in the context of an urban wild duck population largely composed of non-migratory resident individuals, we were able to generate a robust AIV infection and immunity timeline at a population-wide scale, adding to the growing body of literature about these complex dynamics.

### Changes in seroprevalence over time

Over the course of 2011-2014, mean AIV-NP seroprevalence was 27.6%, with some variation between seasons. Sampling over these years typically occurred during the late fall and early winter, during and after typical peak AIV circulation in the region. These data serve to establish a baseline for AIV immunity in this population prior to the incursion of H5N1 into the region. A variety of factors could affect variation in seroprevalence between years, including year-to-year variations among circulating AIV subtypes/strains and population age structure and exposure history. AIV prevalence can follow cyclic patterns, with increased prevalence every several years [57–60]. Prevalence was noticeably higher in 2011-2012, however, these individuals were captured by bait-trapping, which may overestimate true AIV prevalence [61], while ducks were captured by hand in the other years, possibly contributing to this difference.

As expected, AIV seroprevalence increased greatly in the population following the arrival of H5N1 in November 2021, when nearly every duck sampled in January 2022 was seropositive. There was nearly homogenous seroprevalence across all sampled sites and the same banded ducks were observed using multiple urban waterbodies in the area. Given this, along with the extremely high proportion of ducks banded locally only ever being encountered in the same area, we consider the ducks in this urban region at this time to represent a single population. Therefore, there was a population-wide infection event after the arrival of the virus from Eurasia. As AIV-infected waterfowl often exhibit reduced local movements, the large number of infected individuals shedding virus into the local environment could have increased the infection rate at this time, facilitating proficient spread throughout the population to cause near homogenous seroprevalence [9,62]. It is possible that the two seronegative individuals sampled in winter-spring 2022, a mallard and an American wigeon, had moved into the area from elsewhere and were therefore not present at the time of the population-wide infection that occurred roughly a month and a half prior to their sampling. Alternatively, although we believe less likely due to an increase of the number of ducks present in February compared to January, they may have been present in the area but were not infected or were infected but did not generate a detectable antibody response.

Using repeated sampling of this population for roughly 16 months we found that AIV-NP and H5-specific seroprevalence changed greatly in the months following the H5N1 incursion. Individuals were seropositive for anti-NP antibodies for roughly twice as long as H5-specific antibodies, with the population being H5-seronegative roughly 6 months after the incursion. It is well known that waterfowl, and this duck population specifically [38,39], are frequently infected by LPAIVs, which likely explains the longer period of elevated anti-NP antibodies that are boosted with each subsequent infection. Dabbling ducks using human-dominated landscapes in Atlantic Canada show notably high survival rates and strong annual site fidelity to wintering areas [63], leading to a population with an older age distribution than usual. This older age structure may have contributed to the higher seroprevalence overall and over time, as antibody levels are elevated and persist longer in older individuals [28,64,65]. With low seroprevalence of H5-specific antibodies from 2012-2014, assumed to be due to occasional circulation of LPAI H5 viruses, most individuals were likely infected for the first time with an H5 virus, explaining the shorter period in which these specific antibodies persisted.

A slight increase in AIV-NP seropositivity was observed between weeks 32 and 37 (late July to early September 2022), when several individuals tested positive for LPAIVs. However, in contrast to the initial HPAIV H5N1 arrival, the circulating LPAIVs did not infect the whole population but seem to have provided a boost in AIV immunity for some individuals. As these positive ducks were not previously banded, we do not know whether they were migratory or resident individuals. However, due to the time at which this infection occurred and the HA subtypes detected, we suspect that these viruses entered into the population via migratory waterfowl, and not through infection with AIVs that persisted in environmental reservoirs [66,67]. A second appreciable change in AIV-NP and H5-specific seropositivity can be seen between weeks 58 and 59 (February 2023) that can likely be attributed to small variations between locations and the circulation of LPAIVs at these locations. Ducks were caught at Bowring Park on week 58, and both Bowring Park and Quidi Vidi on week 59, with the latter location having a much larger population of birds at that time of the year, including several hundred gulls that include some originating from the Arctic and Europe. This was also shortly after a harsh winter storm, when several species of diving ducks (greater and lesser scaup (*Aythya marila*, *A. affinis*), tufted ducks (*A. fuligula*), ring-necked ducks (*A. collaris*), and red-breasted and common mergansers (*Mergus serrator*, *M. merganser*)) that would normally be using coastal marine habitat at this time of the year [68] took shelter at the lake, which may have contributed to this change.

On 7 February 2023, a single northern pintail (1/7, 14.3%) was seropositive for both AIV-NP and H5-specific antibodies and a week later AIV RNA was detected in the population, signalling the likely circulation of an H5 virus in the region once again. Based on the declining H5-specific antibody seroprevalence since the original population-wide infection event and the lack of H5 viral RNA detected, H5-specific seropositivity was presumably very low until the time at which this individual tested positive in February 2023. The increased detections of H5-specific antibodies in the following weeks, though in a low number of individuals, signaled low level circulation of an H5 virus in the population during this period.

Three weeks later, in March 2023, H5 viral RNA was again detected, and all birds sampled were AIV-NP seropositive. This included a Eurasian wigeon that likely arrived in February 2023 based on sightings of several flocks at this time, and a lesser scaup that likely came into the urban area with other diving ducks to shelter from the harsh winter storm. Only one of these individuals was seropositive for H5-specific antibodies at this time, with the differences likely due to a quicker memory response, an anamnestic response, for AIV-NP antibodies due to frequent infection [27,28,30]. By the end of April 2023, all individuals sampled were seropositive for both AIV-NP and H5-specific antibodies, indicating that an H5 virus had again caused population-wide infection. Although the population was fully infected a year prior, and roughly half of the population still had elevated AIV-NP antibody levels when an H5 virus re-appeared, this was not sufficient to protect against infection. The HPAI H5N1 virus that appeared in March 2023 was different than the original virus from the winter of 2022 and represented a new incursion in the region (Wight et al., unpublished data) with the second infection event of the duck population likely occurring sometime in February 2023. While it is unknown how this new H5N1 entered into the population, the mixing of diving ducks with the urban population may have served as a route of transmission. Hundreds of gulls originating from Arctic, European, and mainland North American breeding populations congregate at Quidi Vidi lake each winter, which is the same location as the first detections of the new H5N1 in the urban duck population (Supplementary Data). Previous work from our group has identified gulls as important vectors by which Eurasian clade AIVs enter into North America [69–71], thereby serving as an alternative explanation for how the new H5N1 may have entered into the urban duck population.

The timing of sampling happened to coincide with the start of low-level circulation of this new H5N1 virus in the population before it subsequently resulted in a second population-wide infection event. This re-infection of the population just over a year later could have been due to a variety of factors. The higher virulence and infectivity of HPAIVs compared to LPAIVs likely played an important role in the original and subsequent population-wide infection events [11,13,72–74]. Waning immunity that did not provide protection from re-infection, escape from the immune system due to low cross-reactive antibodies owing to differences between the two viruses, and delay in memory responses that would allow viral infection to occur, may have also played a role.

The majority of AIV homologous and heterologous challenge studies have been performed along short time scales, usually a matter of weeks, but a few recent studies have evaluated viral shedding duration coupled with serological responses over much longer time scales. An infection study by Shriner et al. [28] found that snow geese (*Anser caerulescens*) infected with an H4N6 virus exhibited minimal viral shedding and antibody levels increased to a level considered seropositive at 7 days post infection (dpi), peaked at 10 dpi, then waned over the next several months and reached undetectable levels one year after the infection. Other studies on wild gulls have found detectable AIV antibody levels for up to a year, demonstrating that responses in some individuals can be long-lived, likely boosted through re-exposure [29,65]. Long-lived responses were also found in a homo- and heterologous challenge study of captive mallards with several different AIVs, detecting AIV-specific antibodies in some individuals more than a year later, showing long term antibody persistence is possible even without boost by re-infection/re-exposure [27]. However, due to the lack of homo- and heterologous challenge studies with HPAIVs in waterfowl along timescales of months to years, periods that are relevant to timing of bird migration, it is currently unknown how long HPAIV-specific antibodies remain elevated and how long individuals are protected from subsequent re-infection.

Migratory individuals with lower AIV seroprevalence that arrived in the region as well as AIV-naïve ducks born in the summer of 2022 may have also contributed to the spread of the 2023 H5N1 virus among the population. The success at which these viruses spread throughout the population during both events may have also been aided by the time of year in which this occurred. For ducks in the Northern Hemisphere, AIV infections typically peak in the fall of each year [3] and the two H5N1 infection events both occurred in early winter. During this time, elevated levels of antibodies present during the peak fall infection period would be waning and low energy stores due to reduced food availability and colder conditions may have made birds more susceptible to infection [22,50,75–77]. Additionally, increased density of birds due to frozen waters may have increased the likelihood of infection during this time. Following the detection of the newly introduced lineage H5N1 in early 2023, several mute swans (*Cygnus olor*) and a number of American black ducks in St. John’s were reported dead, while there was no documented mortality of waterfowl in the region when H5N1 initially infected the population in late-2021/early-2022 [37].

### Relationship between viral load and antibody levels

Extending beyond population-scale seropositivity, we used S/N ratios as a measure of antibody levels along with AIV RNA load (based on Ct values) to investigate patterns of immune response over the course of infection on an individual scale. As individuals progress along the course of infection and transition from the viremic to immunologic phase, viral RNA decreases while antibody levels begin to increase. With substantial variations by species, virus, and body condition, previous work has found that AIV shedding often peaks between 1-8 dpi and lasts for 5-11 days, although some individuals may shed virus for several weeks [22–25,32,50,78,79]. The period of viral shedding has also been found to decrease with more frequent infections [64], and therefore there is a very small window in which ducks can be caught and documented with an active infection [26].

After classifying individuals based on infection status, we found that recently infected individuals had a range of antibody levels that matched individuals sampled closer to the date of the population-wide infection and having higher levels than those infected many months prior (**Fig 4**). This pattern was further clarified by examining individuals that were infected one, three, and six months prior, with antibody levels in each of these groups declining significantly over time. As these are wild birds, many of which were captured for the first time when sampled, we are unable to determine their previous infection history. In contrast to recently infected individuals, no pattern was observed with the timeline of infection and epidemiology or the antibody levels for naïve or waned birds. Although age of each bird was recorded, there appeared to be no differences in antibody levels for seronegative individuals, i.e., seronegative HY birds did not have lower S/N ratios than AHY, SY, or ASY individuals, agreeing with previous observations [50]. We did not pursue analyses by age structure or between sexes due to the limited sample size of each group for each sampling event. Maternal antibodies passed into the egg may have provided some protection to HY birds, at least for a short period, with one study detecting AIV antibodies for up to 17 days in mallard ducklings post hatching [80]. Although we currently do not know the prevalence at which maternal antibodies are found in this population, the expected short persistence of such maternal antibodies is unlikely to contribute to substantial protection within the population overall.

### Immune responses in currently infected individuals

Although only nine ducks were sampled while actively shedding AIV, there was a negative relationship trend for antibody levels versus viral RNA load. This pattern was further supported by inclusion of additional data from seabirds in order to provide a larger dataset of currently infected individuals and the relationship of increasing antibody levels with decreasing viral RNA load is clear (**Fig 5B**). This is an expected immunological response and shows that innate and memory immune mechanisms are quickly responding by generating antibodies as individuals are clearing the infection and leaving the viral shedding phase [30]. This also allowed us to infer the phase of infection at an individual level. For birds recently infected with AIV and therefore already having an elevated antibody titre, their antibody level at the time of sampling may also be confounded by the seroconversion occurring from their ongoing active infection [9]. We are unable to determine previous infection history of each individual as we are interpreting this relationship as a whole population, but factors such as age, infection history, as well as species-level differences would be expected to affect antibody levels on an individual scale [28,29,50,65,78].

### Changes in serology of two recaptured ducks

Based on HPAIV prevalence and epidemiology over the course of this study, it is unlikely that either of the recaptured northern pintails were infected with HPAIV between the two sampling points. However, the male (1196-13442) was likely infected by an LPAIV at some point between the two sampling events as its antibody levels hardly changed between sampling events, roughly one year apart, and it did not have elevated H5-specific antibodies (**Fig 6**, Supplementary Data). Captive infection studies have shown that AIV antibodies persist for several months, however antibodies have not been found to remain elevated to this degree for over a year, even in older individuals [24,25,28,29,32,65]. In contrast, the second individual (female, 1196-13448) seems unlikely to have been infected with an LPAIV between sampling events and, despite LPAIVs being detected at Bowring Park the same day as its original sampling, it was seronegative (**Fig 6**). In light of previous findings from captive infection studies, even if this individual became infected soon after initial sampling, elevated antibody levels would likely still have been detected when resampled approximately six months later. Although these data come from only two individuals, using the combined AIV prevalence, seroprevalence, and epidemiological approach helps add to our understanding of AIV dynamics in wild populations. Efforts to resample individuals multiple times from locations with known AIV dynamics and population movements would be of substantial interest for future studies to evaluate changes in seroprevalence more thoroughly, particularly on an individual basis, and how this contributes to population level immunity [30,81].

## Conclusions

In this study we used a combined approach of screening wild birds for both active AIV infection and for serum antibodies to detect past infections. These data, coupled with known bird movements and epidemiology of the ongoing HPAIV outbreak, allowed a thorough investigation of infection and immunological responses in an urban duck population over a period of 16 months following the arrival of HPAIV from Europe. This study was possible due to the known bird movement and AIV history prior to the arrival of HPAIV for this primarily resident duck population that could be repeatedly sampled, and adds to the growing body of literature highlighting the need for more studies of AIV infection and immunity patterns in wild birds [9,30]. Wildlife surveillance of infectious diseases is a critical aspect of preparedness within a One Health framework and is particularly important with respect to HPAIV, which is a multispecies pathogen with impacts far beyond the poultry industry.

## Acknowledgments

The authors would like to thank all additional individuals who contributed to sampling efforts including A. Bond (2014), S. Duffy (2013), K. d’Entremont (2022), D. Fifield (2011), D. Fife (2023), S. Avery-Gomm (2013), A. Hedd (2023), R. Hoeg (2022), B. Kelly (2022), A. Kroyer (2014), B. Montevecchi (2022), H. Munro (2014), D. Pirie-Hay (2012-2014), P. Ryan (2012, 2013, 2023), K. Studholme (2022), B. Turner (2023), M. Wallace (2023), S. Wallace (2023), and C. Ward (2022).

## Author Contributions

Conceptualization: JW, GJR, ASL

Data Curation: JW, IR, WX

Formal Analysis: JW, GJR

Funding Acquisition: GJR, KH, ASL

Investigation: JW, IR, HLW, JTC, SR, GJR, RSR, WX, DZ, TNA, YB, KEH, ASL

Methodology: JW, IR, HLW, GJR, TA, WX, DZ, YB, ASL

Project Administration: JW, ASL

Resources: JW, IR, HLW, JTC, SR, GJR, TNA, YB, KEH, ASL

Software: JW

Supervision: ASL

Validation: JW

Visualization: JW

Writing – Original Draft Preparation: JW

Writing – Review & Editing: JW, IR, HLW, JTC, SR, GJR, RSR, WX, DZ, TNA, YB, KEH, ASL

## Figure Legends

**Table S1.** Detailed records for samples from ducks that were used in this study. Samples from the 2011-2012 period were no longer available, therefore the previously published data [38] were used and are presented as only anti-NP antibody-positive or -negative.

